# Numerous cultivated and uncultivated viruses encode ribosomal proteins

**DOI:** 10.1101/174177

**Authors:** Carolina M. Mizuno, Charlotte Guyomar, Simon Roux, Régis Lavigne, Francisco Rodriguez-Valera, Matthew B. Sullivan, Reynald Gillet, Patrick Forterre, Mart Krupovic

## Text

Viruses modulate ecosystems by directly altering host metabolisms through auxiliary metabolic genes, which are obtained through random ‘sampling’ of the host genome and rise to fixation, presumably through improved viral fitness by alleviating key metabolic bottlenecks during infection. Conspicuously, however, viral genomes are not known to encode the core components of translation machinery, such as ribosomal proteins (RPs), though genes for RPs S1 and S21 have been detected in viral metagenomes^1,2^. Here we augment this little-noticed observation using available reference genomes, global-scale viral metagenomic datasets, and functional assays for select proteins. We identify 15 different RPs across diverse viral genomes arising from cultivated viral isolates (5 RPs in 16 genomes) and metagenome-assembled viruses (14 RPs in 1,403 uncultivated virus genomes). Among these, S21 and L7/L12 are the most common, and functional assays show that both proteins are incorporated into 70S ribosomes when expressed in *Escherichia coli*, indicating that they might modulate protein translation during infection. Ecological distributions of virus-encoded RPs suggest ecosystem-specific virus adaptations, whereby aquatic viruses appear to selectively incorporate genes for S21, L31 and L33, whereas S6, S9, S15 and S30AE genes are enriched among viruses infecting animal-associated bacteria. Finally, the fact that viruses tend to encode dynamic RPs, suggests that the viral proteins likely replace cellular versions in host ribosomes, likely enabling takeover of host translational machinery.

During billions of years of co-evolution with their hosts, viruses have evolved numerous strategies ensuring their successful propagation, including tinkering with various metabolic pathways and subversion of key cellular biosynthetic machineries. For example, ocean viruses that infect cyanobacteria (cyanophages) commonly encode core photosynthetic reaction center proteins, which serve to maintain the complex photosynthetic machinery during infection^3^. These and other ocean viruses can similarly manipulate their host’s ability to uptake phosphate^4^, as well as cycle nitrogen^5,6^ and sulfur^7,8^ – the fundamental building blocks of ocean life. Complementarily, viruses employ a diverse array of host take-over strategies to (i) fight off host defenses by encoding anti-restriction-modification or anti-CRISPR genes^9^, and (ii) control transcription by encoding sigma factors or polymerases themselves ^10^. Conspicuously not yet observed in viral genomes, however, are the RPs. Indeed, even the giant mimiviruses, which are known to carry genes for a range of aminoacyl-tRNA synthetases^11,12^, do not encode proteins directly participating in the formation of the ribosomes. It is this feature – ribosome-encoding or not – which is now proposed to separate cellular life from viruses^13,14^.

The first crossing of this line appeared when previous analysis of ‘cleaned’ viral metagenomes suggested that viral genomes might encode ribosomal proteins after all – specifically for S1 and S21 (REF 1,2). While intriguing, these observations went largely unnoticed, likely because they were based on short assemblies lacking genome context. To systematically investigate this, we first searched available reference genomes of cultivated viruses for genes encoding RPs. Of 116 ribosomal protein domains (Table S1) that seeded our searches, 5 were identified across 16 viral genomes (Table 1). The genes were generally embedded within variable genomic contexts, even for homologous ribosomal protein genes (Figure S1).

**Table 1.** Ribosomal protein domains found in cultivated viruses.

| Domain | Name (Family) | Protein<br>accession,<br>length (aa) | Coverage<br>(%) | Identity<br>(%) | Probability<br>(%) | E-value |
| --- | --- | --- | --- | --- | --- | --- |
| Ribosomal_S30 | Finkel-Biskis-Reilly murine sarcoma virus (*R*) | NP_598374, 133 | 85 | 86 | 99.92 | 2.2E-26 |
| Ribosomal_S21 | Pelagibacter phage HTVC008M (*M*) | AGE60443, 67 | 59 | 46 | 99.81 | 2.7E-19 |
| Ribosomal_L9_N | Mycobacterium phage 32HC (*S*) | AHJ86298, 86 | 33 | 40 | 98.32 | 4.3E-07 |
| Ribosomal_L12 | Dinoroseobacter phage DFL12phi1 (*P*) | AHX01035, 106 | 74 | 32 | 99.9 | 1.0E-23 |
|  | Erwinia phage Ea9-2 (*P*) | AHI60108, 724 | 9 | 32 | 96.87 | 6.3E-03 |
|  | Ralstonia phage RSB3 DNA (*P*) | BAN92321, 98 | 59 | 32 | 99.77 | 2.2E-18 |
|  | Roseophage DSS3P2 (*P*) | ACL81275, 107 | 62 | 28 | 99.44 | 3.7E-13 |
|  | Salmonella phage FSL SP-058 (*P*) | AGF88397, 418 | 16 | 34 | 96.05 | 1.8E-01 |
|  | Salmonella phage FSL SP-076 (*P*) | AGF88198, 418 | 15 | 36 | 96.21 | 2.6E-02 |
|  | Sulfitobacter phage phiCB2047-B (*P*) | AGH07436, 126 | 25 | 47 | 97.06 | 1.8E-03 |
| Ribosomal_S30AE | Cronobacter phage vB CsaM GAP32 (*M*) | AFC21633, 111 | 71 | 34 | 99.96 | 4.2E-28 |
|  | Enterobacteria phage vB EcoM-FV3 (*M*) | AEZ65272, 105 | 74 | 35 | 99.92 | 1.5E-23 |
|  | Escherichia coli bacteriophage rv5 (*M*) | ABI79209, 105 | 74 | 33 | 99.96 | 1.3E-27 |
|  | Escherichia phage 2 JES-2013 (*M*) | AGM12525, 105 | 74 | 32 | 99.96 | 3.0E-28 |
|  | Escherichia coli O157 typing phage 14 (*M*) | AKE47110, 105 | 74 | 33 | 99.96 | 3.4E-28 |
|  | Escherichia phage vB EcoM FFH2 (*M*) | AEZ65272, 105 | 74 | 35 | 99.93 | 7.9E-24 |

\**R*: Retroviridae, *M*: Myoviridae, *S*: Siphoviridae, *P*: Podoviridae

We identified a ribosomal protein S30 domain, a component of the small 40S ribosomal subunit^15^, in the eukaryotic virus, Finkel-Biskis-Reilly murine sarcoma virus (FBR-MuSV), a member of the family *Retroviridae*. This domain was part of the *fau* gene fused to an N-terminal ubiquitin-like domain (Figure S2a). Interestingly, FBR-MuSV has acquired the cDNA copy of *fau* in inverse orientation, and production of the antisense RNA suppresses expression of endogenous *fau* mRNA, which leads to apoptosis inhibition and induces tumorigenesis^15,16^. Although the viral protein is not translated^15^, the antisense transcript affects the production of the cellular *fau*^16^ and thus might have an indirect effect on the ribosome biogenesis.

The remainder of the virus-encoded ribosomal proteins – S21, L9, L7/L12 and S30AE – was found in bacterial viruses (bacteriophages) infecting proteobacterial (from 3 different classes) and mycobacterial (phylum Actinobacteria) hosts (Table 1). The S21 homolog was identified in pelagiphage HTVC008M, a myovirus. S21 is a conserved component of the bacterial 30S ribosomal subunit (Figure 1a) required for the initiation of polypeptide synthesis and mediates the base-pairing reaction between mRNA and 16S rRNA^17^. The viral protein was most similar (54% identity over the protein length) to the corresponding protein of its host, *Pelagibacter ubique* (Figure 1b), an abundant member of the SAR11 clade (class Alphaproteobacteria), which is considered to represent one of the most numerous bacterial groups worldwide^18^. Maximum likelihood phylogenetic analysis strongly suggests that the phage gene was horizontally acquired from the *Pelagibacter* host (Figure 1c).

**Figure 1.**
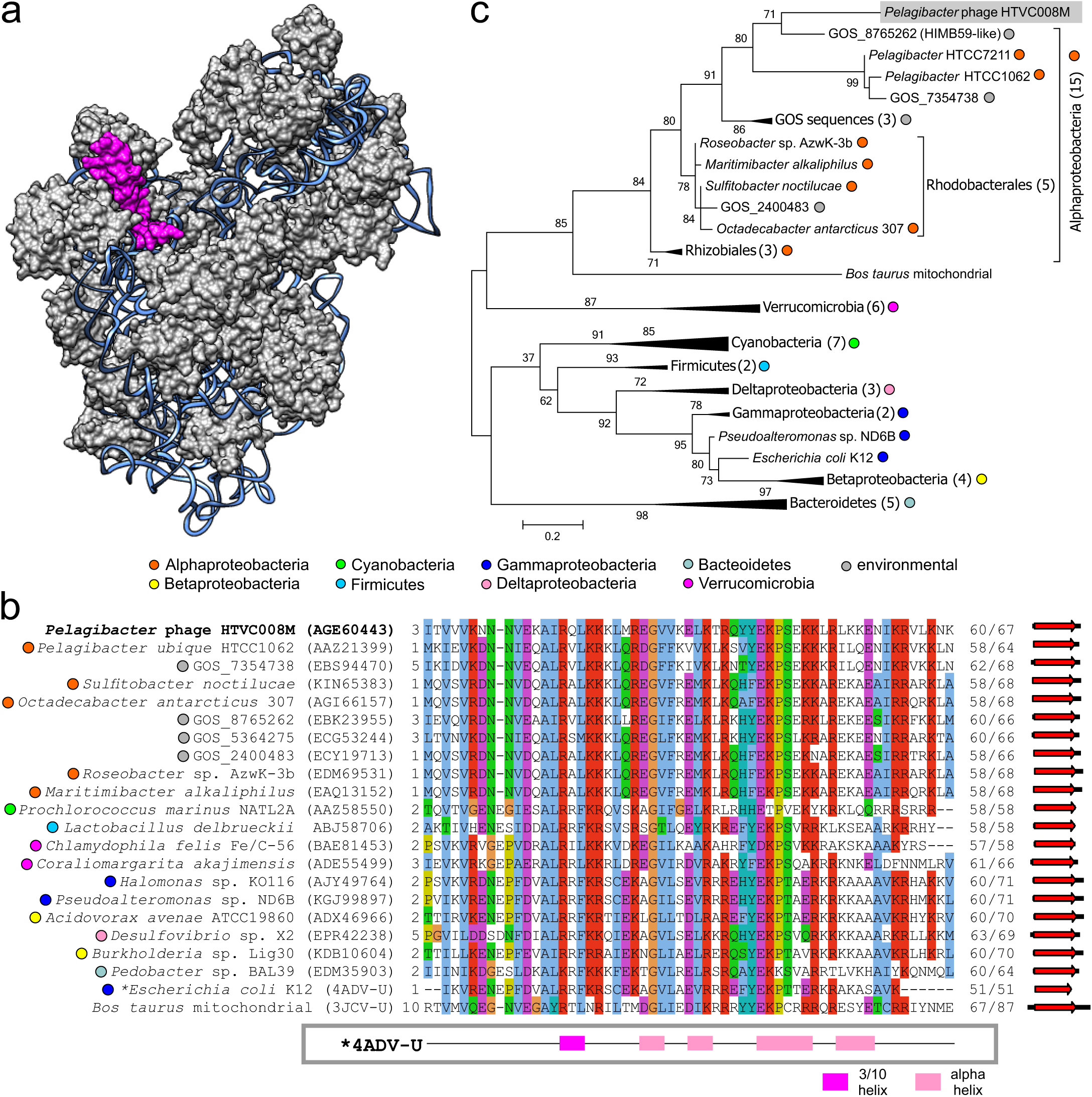
Virus-encoded ribosomal protein S21. **a)** Structure of the *Escherichia coli* 30S ribosomal subunit (PDB id: 4ADV). 16S ribosomal RNA is shown as blue ribbon. S21 ribosomal protein is highlighted in pink. **b)** Alignment of the ribosomal protein S21 encoded by pelagiphage HTVC008M with homologs from representatives of distinct bacterial taxa and environmental sequences obtained from the Global Ocean Sampling (GOS) dataset. **c)** Phylogenetic tree of ribosomal protein S21. Taxonomic affiliations are represented by colored circles (see legend).

Ribosomal protein L9 was identified in *Mycobacterium* phage 32HC, a siphovirus. L9 binds to the 23S rRNA and is a component of the large 50S ribosome subunit (Figure S3A). The protein is involved in translation fidelity and is required to suppress bypassing, frameshifting, and stop codon “hopping”^19^. L9 has a highly conserved architecture consisting of two widely spaced globular domains connected by an elongated α-helix^20^. While the viral L9 homolog contains the N-terminal globular domain and part of the α-helical spacer, the C-terminal part has been apparently non-homologously replaced with sequence that lacks known function (Figure S2b).

The next ribosomal protein encoded in sequenced viral genomes was L7/L12, which was found in 7 phages infecting proteobacteria from 3 different classes (Table 1). L7 is equivalent to L12 except for the acetylation at the N-terminus, hence the two proteins are often collectively referred to as L7/L12^21^. The L7/L12 proteins participate in the formation of the so-called L7/L12 stalk, a clearly defined morphological feature in the *E. coli* 50S ribosomal subunit, which besides L7/L12, contains ribosomal proteins L10 and L11 as well as the L10- and L11-binding region of the 23S rRNA^21^ (Figure S3A). The phage-encoded L7/L12 domains are similar (∼50%) to *bona fide* cellular ribosomal homologs, as well as conserved residues involved in interaction with L11 and elongation factors EF-G and EF-Tu (Figure S4a). Although in some phages (e.g., *Ralstonia* phage RSB3), the L7/L12 domain spans the entire protein, it was more common that these domains were variably positioned within much larger polypeptides (up to 724 aa-long; Figure S4b). Interestingly, searches seeded with sequences flanking the L7/L12 domain in phage proteins resulted in identification of multiple phage homologs which specifically lack the L7/L12 domain (Figure S5). For example, proteins encoded by *Salmonella* phages FSL_SP-058 and FSL_SP-076 contain the L7/L12 domains, whereas homologous protein from *Escherichia* phage Pollock lacks this domain, despite conservation of the upstream and downstream regions (Figure S5). Furthermore, in different phage genomes, L7/L12 proteins were encoded within widely different genomic contexts (Figure S1). These observations suggest that L7/L12 domain has been acquired by different phages on multiple, independent occasions, with some of these genes possibly being fixed in the phage genomes.

The last ribosomal protein encoded in sequenced viral genomes were S30AE domain-containing proteins, which were encoded by 7 phages infecting *Cronobacter* and *E. coli* (six closely related phages with 92-97% average nucleotide identity) (Figure S6). S30AE proteins are expressed during stasis and under unfavorable growth conditions. S30AE proteins binds ribosomes to stabilize 100S dimers that inhibit translation to enable cells to control translational activity without costly alteration of the ribosomal pool^22^. Multiple sequence alignment shows high conservation of the viral and cellular S30AE homologs (Figure S7), suggesting that the gene transfer has occurred in a relatively recent past. In the S30AE phylogeny, homologs from *E. coli* phages cluster amidst gammaproteobacterial sequences. By contrast, the more divergent protein encoded by *Cronobacter* phage clusters with sequences from members of the phylum Firmicutes, though this association is confounded by potential long-branch-attraction artifact (Figure S8).

To place these findings of cultivated virus-encoded RPs into broader ecological context, we searched 424,225 viral contigs from two global viral metagenomic datasets^8,23^ for putative RPs using the same 106 sequence profiles (see Materials and Methods). Overall, 14 putative ribosomal protein genes were identified across 1,403 contigs (Figure 2, Table S2, Figure S3B). S21, L7/L12 and S30AE, which were found in cultivated phages, were also detected in uncultivated phages, with S21 homologs dominating (88%) the pool of RPs detected (Figure 2, Table S2). While found in only one cultivated phage (see above), maximum likelihood phylogeny and genome context comparison using these metagenomic data suggested that at least 7 virus-host exchanges of S21 protein-coding genes have occurred, and across multiple bacterial phyla (Figure 2, Figure S9). Notably, S21-encoding viruses were almost exclusively from aquatic samples (90% of S21s detected). Such repeated transfers and enrichment in aquatic samples suggest that virus-encoded S21 proteins likely can provide a direct fitness benefit to aquatic bacteriophages. By contrast, L7/L12 and S30AE were found across a broad range of samples (Figures 2, S6, S10), suggesting that their repeated acquisition could be beneficial in multiple types of conditions and hosts.

**Figure 2.**
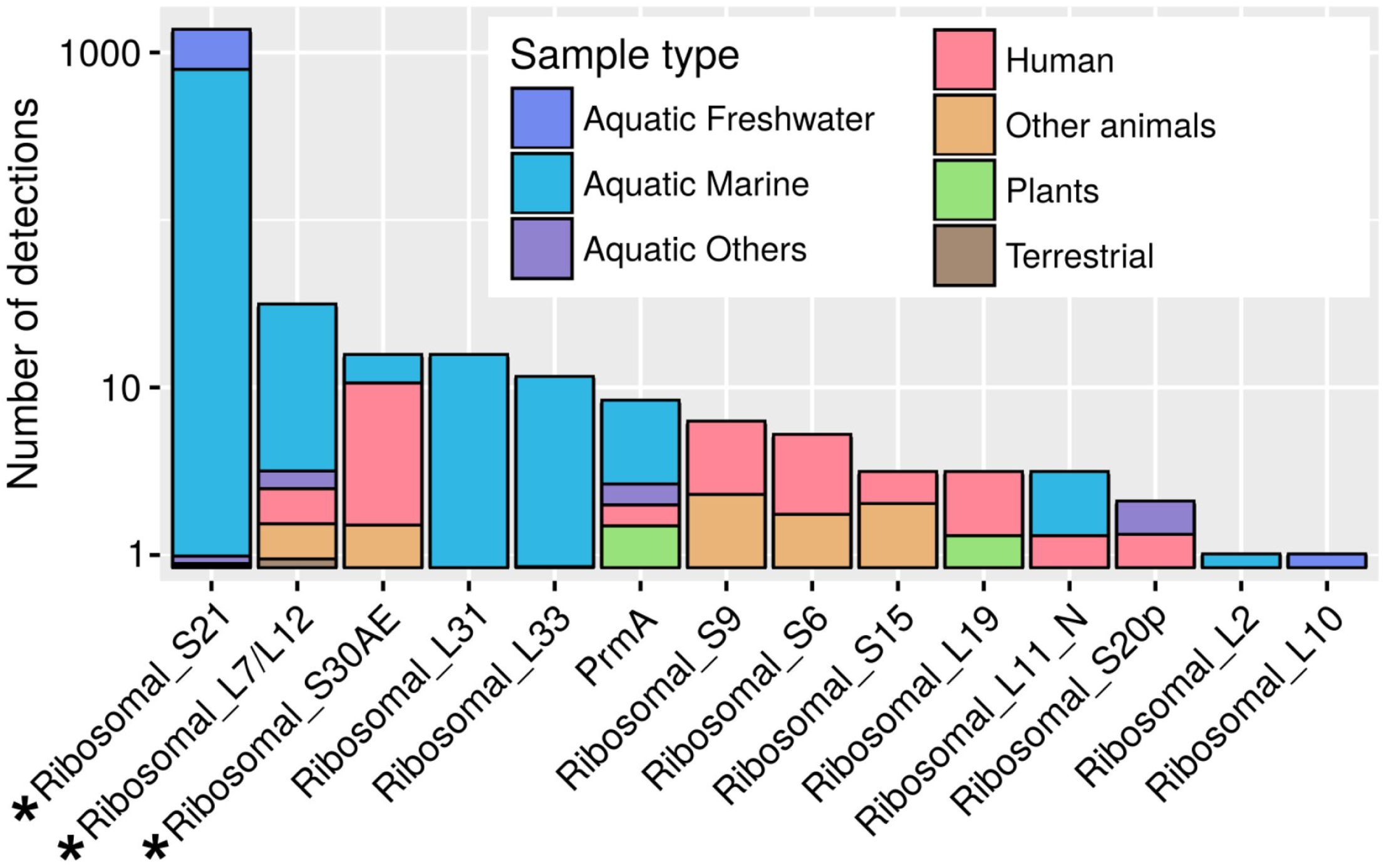
Detection of ribosomal proteins in uncultivated viral genomes (assembled from metagenomes). For each ribosomal protein detected, the total number of detection is shown on the y-axis (log_10_ scale), and the bar is colored according to the type of samples in which this protein was detected (the sizes of the colored parts are proportional to the number of detections made in each type of samples). Ribosomal proteins also identified in cultivated viruses are identified with stars.

Additionally, however, we identified another 11 RPs in uncultivated viruses that were not identified in the isolate genomes (Table S2). Notable among these, due to being commonly (>10 viral contigs) detected, are L31 and L33. Although the biological function of L33 remains obscure ^24^, it appears to contact tRNAs in the ribosomal E(exit)-site^25^, whereas L31, similar to S30AE, plays a role in 100S formation, 70S association, and translation^26^. As in the case of S21, viral contigs encoding L31 or L33 were almost exclusively detected in aquatic environments (Figure 2). Maximum likelihood phylogenies and genome context comparisons highlighted a consistent pattern of at least 2 independent events of virus-host transfers involving viruses infecting different bacterial phyla (Figures S10 and S11).

Thus, at this point, there is an emerging picture that viruses might randomly sample host DNA, including ribosomal protein genes, and that in some cases these might become fixed in viral genomes. Most (>99%) of the viruses contained only a single ribosomal protein gene (exception: 9 uncultivated viral contigs contained 2; Figure S12), which is clearly not enough for viruses to build functional ribosomes on their own. Presumably, these viruses are merely tweaking ribosomal functioning in their hosts – just as observed for auxiliary metabolic genes whereby viruses do not encode complete pathways, but instead only select genes critical for the takeover and/or reprogramming of the host cell^6,7,27^.

Presence of ribosomal protein genes in viral genomes raises a question of what their functions in the course of the infection cycle might be and how do viruses benefit from carrying such genes. The S30-encoding gene increases the transformation capacity of FBR-MuSV *in vitro* by twofold, providing clear fitness advantage to the virus^15^. It is conceivable that homologs of other ribosomal proteins might be also beneficial for the bacteriophages that encode them. For instance, it is known that S21 is necessary during translation initiation step and in the absence of S21, ribosomes are incapable of binding natural mRNAs^17^. Thus, phage-encoded S21 might compete with and replace the cellular S21, forcing preferential translation of viral transcripts. Similarly, viral L7/L12 domain proteins might provide interfaces for virus-specific translation factors. Protein L9 is required for translational fidelity and is involved in suppression of frameshifting. In many members of *Caudovirales* production of certain tail components is dependent on programmed translational frameshifting ^28^ and viral copy of L9 might help to achieve optimal frameshifting in these genes. It has been demonstrated that stalling of phage protein synthesis is one of the major defense strategies in Bacteroidetes^29^. Thus, viral homologs of S30AE and L31 might compete with the cellular homologs and prevent formation of ribosome dimers, thereby releasing translation inhibition and ensuring that phage transcripts are efficiently translated.

Given what seemed to be reasonable explanations for why viruses might benefit from encoding such genes, we next investigated whether virus-encoded ribosomal protein genes appeared functional. Thus, we calculated the ratio of nonsynonymous polymorphisms per non-synonymous site (pN) to the number of synonymous polymorphisms per synonymous site (pS). This ratio can be used to infer whether genes are evolving neutrally (pN/pS=1) or positively (pN/pS>1) away from the original function, or whether such substitutions are largely not tolerated due to purifying selection (pN/pS<1) that would suggest the gene was functional. These analyses suggested that the vast majority of the viral-encoded RPs were likely functional as well-sampled genes (>10x coverage, and ≥1 single nucleotide polymorphism, or SNP) had an average pN/pS=0.10, with 84% having a pN/pS≤0.20 (Table S2).

To build on these *in silico* functional assays, we next explored whether the viral proteins are incorporated into ribosomes, by focusing on 3 RPs encoded by cultivated phages and most frequently detected in uncultivated phage genomes (Figure 2). These were pelagiphage-encoded S21, L7/L12 from *Salmonella* phage FSL SP-076, and S30AE from *Escherichia coli* phage rv5. Following moderate and controlled expression of the respective viral proteins, 70S ribosomes were isolated under high-stringency salt conditions (see Materials and Methods) to avoid unspecific association of viral proteins^30^. Judging from the obtained ribosome profiles (Figure 3A) and transmission electron microscopy (Figure 3B), expression of the viral proteins did not affect the 70S stability. All examined samples nearly exclusively contained 70S monoribosomes. Subsequent mass spectrometry (MS) analysis of the 70S ribosomes purified on the sucrose gradients unequivocally showed that S21 and L7/L12 (Figure 3C, Supplementary Table S4 and S5), but not S30AE (Supplementary Table S6), were stably incorporated into the 70S ribosomes when expressed in *E. coli*. Notably, S30AE was detected using MS in the crude cell extracts (Figure 3C, Supplementary Table S7), indicating that lack of its incorporation into ribosomes is not due to poor protein expression, but may rather result from other factors, such as inadequate growth phase, genuine loss of ability to bind to ribosomes or dissociation due to stringent washes with salt during 70S ribosome isolation. Regardless, binding of viral S21 and L7/L12 to ribosomes strongly suggests that these and possibly other viral RPs modulate protein translation during phage infection.

**Figure 3.**
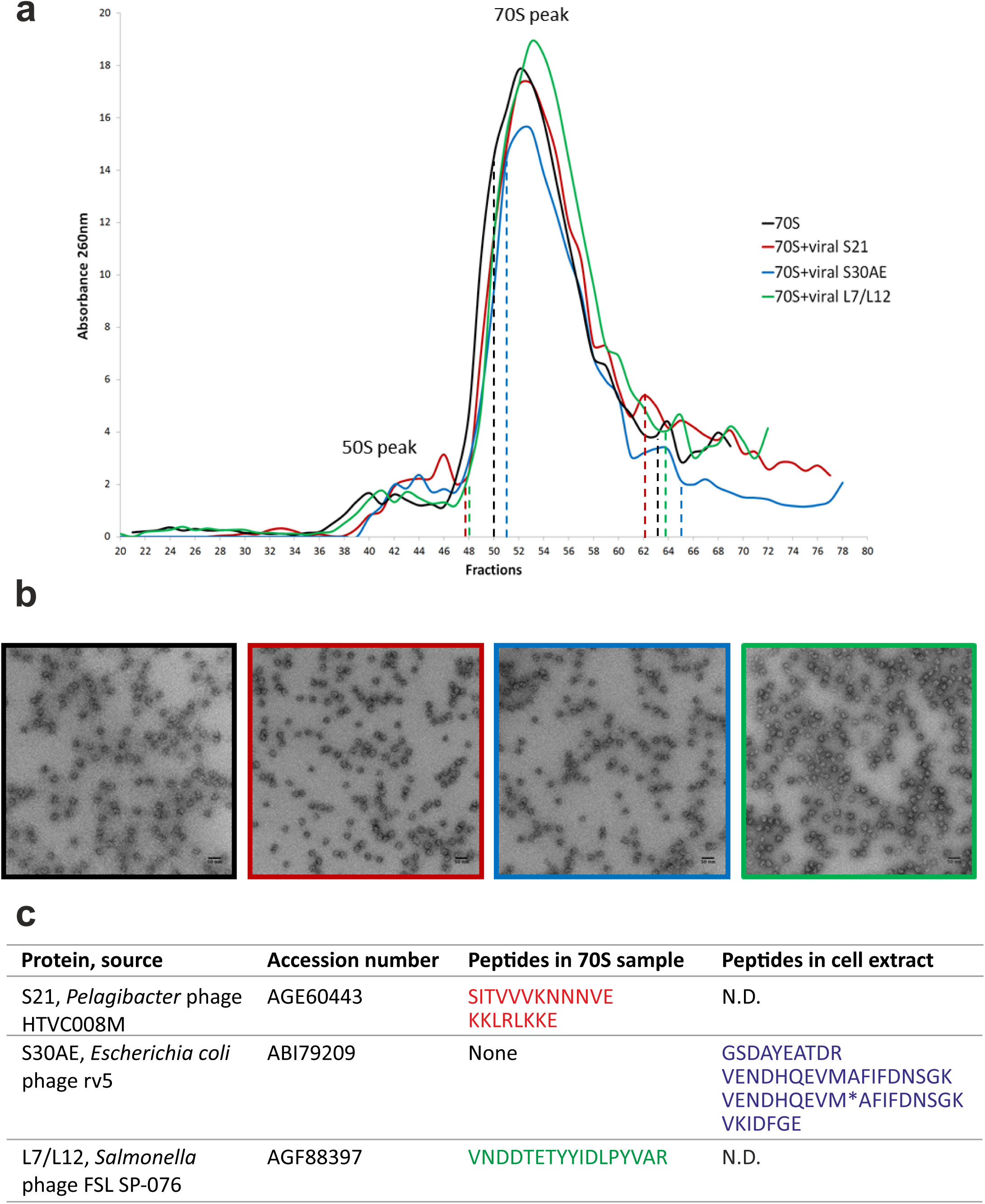
Ribosome analysis of extracts from NM522 *Escherichia coli* cells. **a)** A260 profile of ribosome extracts separated on a sucrose gradient. *Escherichia coli* cells expressing viral S21 (red), S30AE (blue) or L7/L12 (green) were separated through a 10%–50% sucrose gradient and fractionated to be compared to the same strain in the absence of induction (black curve) (see experimental section). The dotted lines indicate the fractions that were pooled and analyzed by mass spectrometry. **b)** The corresponding 70S fractions (dot lines, same color codes) were visualized on electron micrographs of 70S ribosomes negatively stained with 2% uranyl acetate. Scale bars: 50 nm. **c)** S21, S30AE and L7/L12 peptides identified by mass spectrometry in ribosome preparation and crude cell extract, respectively. Asterisk denotes oxidized form of methionine. N.D., not determined.

In summary, this work builds upon prior discoveries of aminoacyl-tRNA synthetase genes encoded by giant viruses (family *Mimiviridae*)^11,12^. Our current work shows that even ribosomal proteins are encoded by numerous cultivated and uncultivated viruses with relatively small genomes, and offers support for them having an evolutionary fitness advantage for viruses during infection. Interestingly, virus-encoded RPs appear to be differentially selected for across environments as aquatic viruses are enriched for S21, L31 and L33, whereas phages of animal-associated bacteria are enriched for S6, S9, S15 and S30AE. Curiously, although ribosomes are highly stable macromolecular assemblies which retain most of their original components during cellular growth and division^31^, some elements (proteins S21, L7/L12, L9, L31 and L33) are highly dynamic, solvent accessible, and among the few proteins that are loosely bound to the ribosome and can be exchanged *in vivo* between ribosomes^31,32^. It is these dynamic ribosomal proteins that are enriched in viruses, which presumably is because they are most suited to homologous replacement during infection and therefore of a functional fitness advantage during phage evolution. Analogously, modulation of photosynthesis as well as nitrogen and sulfur cycles in infected cells hinges on a handful of key proteins captured by viruses from their respective hosts^6,8,33^. More generally, such selective acquisition of key components of the multisubunit assemblies, such as ribosomes and photosystems, or recruitment of central regulators of rate-limiting steps in metabolic pathways appears to be a general strategy employed by viruses to optimize the metabolic state of the infected cells and/or to achieve the takeover of the host.

## METHODS

### Sequence analyses

All viral genomes were downloaded from viral RefSeq database (ftp://ftp.ncbi.nlm.nih.gov/refseq/release/viral/). A hidden Markov model (HMM) profile was downloaded from the PFAM database (http://pfam.xfam.org/) for each domain listed in Table S1. In total, 106 sequence profiles corresponding to distinct ribosomal protein domains were used as seeds to search the proteomes of viruses infecting hosts from the three cellular domains, as well as proteins predicted on viral contigs from two previously published global metagenomic datasets, Global Ocean Virome^8^, and Earth’s Virome^23^, which are available at https://img.jgi.doe.gov/cgi-bin/vr/main.cgi and http://datacommons.cyverse.org/browse/iplant/home/shared/iVirus/GOV. Notably, domain S1, which is repeated 4 to 6 times in the ribosomal protein S1, is not exclusive to RPs as it is common across diverse RNA-binding proteins and fused to non-ribosomal functional motifs (pfam id: PF00575.18). Thus while domain S1 was found in homologs of vaccinia virus interferon inhibitor K3L^34^, which is conserved in chordopoxviruses belonging to 7 different genera, it was not considered further due to potential functional ambiguity. The domains were identified by HHsearch ^35^ with E-value of 1e-5. For isolates, the identified hits were then manually inspected using HHPRED ^35^. All alignments were constructed using PROMALS3D^36^. Maximum likelihood phylogenetic trees were constructed using PhyML^37^ using a WAG substitution model and the proportion of invariable sites estimated from the data. For metagenomic predicted proteins, multiple alignments were built with Muscle^38^ and maximum likelihood phylogenetic trees were computed with FastTree^39^, and displayed with iTol^40^. Genomic comparisons were performed using BLAST with the BLOSUM45 matrix. The ribosomal structure was downloaded from PDB database and visualized using Chimera^41^.

To further confirm the functionality of RPs encoded on uncultivated viral genomes, selective constraint on these auxiliary metabolic genes was evaluated through pN/pS calculation, as in REF. 42. Briefly, synonymous and non-synonymous SNPs were observed in each ribosomal protein gene covered ≥ 10x, and compared to expected ratio of synonymous and non-synonymous SNPs under a neutral evolution model if at least 1 SNP was identified. The interpretation of pN/pS is similar as for dN/dS analyses, with the operation of purifying selection leading to pN/pS values < 1.

### Genetic constructions

The genes encoding for S21 protein from *Pelagibacter* phage HTVC008M (AGE60443), S30AE protein from *Escherichia coli* bacteriophage rv5 and L7/L12 protein from *Salmonella* phage FSL SP-076 (AGF88397) were synthetized by Eurofins Genomics (Ebersberg, Germany). S21 and S30AE genes were cloned into pEX-A2 plasmid and L7/L12 gene into pEX-K4 plasmid. The gene corresponding to S30AE viral protein was digested by BsaI and HindIII and inserted into a pBAD24 vector between NcoI and HindIII restriction sites. The genes corresponding to S21 and L7/L12 viral proteins were cloned into the same vector, using EcoRI and HindIII restriction sites. The pBAD24 plasmid harbors an arabinose dependent promoter, a pBR322 origin and the ampicillin resistance coding sequence.

### Protein expression and cell retrieval

NM522 *Escherichia coli* strain was used for expression of viral S21, S30AE and L7/L12 proteins. The same strain harboring empty pBAD24 was used as a negative control. Overnight pre-cultures were grown in the presence of 1mM of L-arabinose and 100μg/mL of ampicillin. Then the expression was maintained in the cell culture until the end of exponential phase. Once the cultures reached an OD_600nm_ of 1, the cells were centrifuged at 7,000rpm for 7 minutes at 4°C. The cell pellet was then washed into saline water at a concentration of 9g/L of NaCl. A second centrifugation was made and the bacterial pellet was frozen at -80°C.

### 70S Ribosome purification

The cells were resuspended in Buffer 1 (Tris-HCl pH7,5 20mM,MgOAc 50mM,NH_4_Cl 100mM, EDTA 0.5mM and DTT 1mM) and finally lysed using the French Press. The lysate was centrifuged and the supernatant was put above the same volume of high-salt sucrose buffer (Tris-HCl pH7.5 10mM, MgCl_2_ 10mM, NH_4_Cl 500mM, EDTA 0.5mM, certified RNase free sucrose 1.1M and DTT 1mM) in order to wash the ribosomes. After centrifugation at 30,000rpm for 20h at 4°C, the ribosomes form a translucent pellet. The ribosome pellet was washed several times to remove membranes and then resuspended in Buffer 2 (Tris-HCl pH7.5 10mM, MgCl_2_ 10mM, NH_4_Cl 50mM, EDTA 0.5mM and DTT 1mM) on ice. An equivalent of 200OD_260nm_ units of ribosomes were loaded on top of a 10-50% sucrose gradient into polycarbonate tubes. The ultra-centrifugation was performed at 23,000rpm, for 18h at 4°C using SW28 rotor (BECKMAN L-90 ultracentrifuge). The gradient was then fractionated into 500μL aliquots. The OD_260nm_ values were determined for each fraction to locate the 70S absorbance peak. The corresponding fractions were pooled in one volume of buffer 2 and centrifuged at 30,000rpm for 20h at 4°C in order to remove sucrose. The pellet was recovered in buffer 2 and after titration, the ribosomes were ready for mass spectrometry analysis.

### Negative staining

Following ribosome separation, we diluted samples 10 times in Buffer 2 and applied them to freshly glow-discharged 300-mesh collodion/carbon-coated grids. After three washes in this buffer, grids were stained with 2% uranyl acetate for 30 s. The grids were then observed with a Tecnai G2 Sphera transmission electron microscope operating at 200 kV. Images were recorded with a 4000 × 4000 Gatan Ultrascan 4000 CCD camera at a nominal magnification of 50,000×.

### LC-MS/MS proteins identification

#### - Liquid digestion of ribosomal samples

25 μg of ribosomes were digested according to the following protocol: first, 53.5 μl of 50mM ammonium bicarbonate buffer (pH 7,8) was added to the sample to 65 μL total volume. After vortexing 1 minute, tubes were incubated 10 minutes at 80°C and then sonicated for two minutes. Reduction of disulfide bonds step was processed by adding 12.5 μl of 65mM DTT to the sample and was incubated 15 minutes at 37°C after agitation 1 minute. Alkylation of reduced disulfide bonds was realized by adding 135mM iodoacetamide. Microtube was then incubated 15 minutes in the dark at room temperature, under agitation. Finally, proteins were digested overnight at 37°C with 10 μl of either modified endoproteinase glu-c ([0.1 μg/μl.], Promega, Madison, WI) in 50 mM ammonium bicarbonate buffer for S21 (due to high lysine and arginine content in S21) or with modified Trypsine ([0.1 μg/μl.], Promega, Madison, WI) in 50 mM ammonium bicarbonate buffer for S30AE, L7/L12 and control.

#### - Protein Prefractionation and Digestion

Twenty five micrograms of soluble crude protein extracts of *E. coli* were boiled for 10 min with 5 μl of LDS Sample buffer 4X and 2μl of reducing agent (DTT 10X (500mM)). They were then separated on a NuPAGE^®^ Novex^®^ 4-12 % gradient Bis-Tris gel (Invitrogen Corparation, USA) in MES SDS Running Buffer (Invitrogen: 50 mM MES, 50 mM Tris-HCl, 1 % SDS, 1.025Mm EDTA) using Xcell SureLock Mini Cell (Invitrogen).

Gel was stained with EZBlue (Sigma-Aldrich) for 30 min and destained with water over night. Each gel lane was manually cut into 2 slices of approximately the same size in the region of 7kDa-14kDa. The slices were first treated with 50 mM NH_4_HCO_3_ in acetonitrile/water 1:1 (v/v), dehydrated with 100% acetonitrile and rehydrated in 100 mM NH_4_HCO_3_. Next they were washed again with 50 mM NH4HCO3 in acetonitrile/water, 1:1 (v/v) and dehydrated with 100% acetonitrile. The slices were then treated with 65 mM DTT for 15 min at 37 °C, and with 135 mM iodoacetamide in the dark at room temperature. Finally, the samples were washed with 100 mM NH_4_HCO_3_ in acetonitrile/water, 1:1 (v/v), and dehydrated with 100% acetonitrile before being rehydrated in 100 mM NH_4_HCO_3_, washed with 100 mM NH_4_HCO_3_ in acetonitrile/water, 1:1 (v/v) and then dehydrated again with 100% acetonitrile. Proteins were digested overnight at 37 °C with 4 ng/l of modified trypsin (Promega, Madison, WI) in 50 mM NH_4_HCO_3_. Peptides were extracted by incubating the slices first in 80 μl of acetonitrile/ water/trifluoroacetic acid (70/30/0.1; v/v/v) for 20 min, and then in 40 μl of 100% acetonitrile for 5 min and finally in 40 μl of acetonitrile/water/trifluoroacetic acid (70/30/0.1; v/v/v) for 15 min. Supernatants were transferred into fresh tubes and concentrated in a SpeedVac (Thermo Scientific) for 15 min to a final volume of 40 μl.

#### - LC-MS/MS analysis

Shotgun analyses were conducted on a LTQ-Orbitrap XL (ThermoFisher Scientific) mass spectrometer. The MS measurements were done with a nanoflow highperformance liquid chromatography (HPLC) system (Dionex, LC Packings Ultimate 3000) connected to a hybrid LTQ-Orbitrap XL (Thermo Fisher Scientific) equipped with a nanoelectrospray ion source (New Objective). The HPLC system consisted of a solvent degasser nanoflow pump, a thermostated column oven kept at 30 °C, and a thermostated autosampler kept at 8 °C to reduce sample evaporation. Mobile A (99.9% Milli-Q water and 0.1% formic acid (v:v)) and B (99.9% acetonitrile and 0.1% formic acid (v:v)) phases for HPLC were delivered by the Ultimate 3000 nanoflow LC system (Dionex, LC Packings). An aliquot of 10 μL of prepared peptide mixture was loaded onto a trapping precolumn (5 mm × 300 μm i.d., 300 Å pore size, Pepmap C18, 5 μm) over 3 min in 2% buffer B at a flow rate of 25 μL/min. This step was followed by reverse-phase separations at a flow rate of 0.250 μL/min using an analytical column (15 cm × 300 μm i.d., 300 Å pore size, Pepmap C18, 5 μm, Dionex, LC Packings). We ran a gradient from 2–35% buffer B for the first 60 min, 35–60% buffer B from minutes 60–85, and 60–90% buffer B from minutes 85–105. Finally, the column was washed with 90% buffer B for 16 min and with 2% buffer B for 19 min before the next sample was loaded. The peptides were detected by directly eluting them from the HPLC column into the electrospray ion source of the mass spectrometer. An electrospray ionization (ESI) voltage of 1.6 kV was applied to the HPLC buffer using the liquid junction provided by the nanoelectrospray ion source, and the ion transfer tube temperature was set to 200 °C. The LTQ-Orbitrap XL instrument was operated in its data-dependent mode by automatically switching between full survey scan MS and consecutive MS/MS acquisitions. Survey full scan MS spectra (mass range 400-2000) were acquired in the Orbitrap section of the instrument with a resolution of r = 60 000 at m/z 400; ion injection times were calculated for each spectrum to allow for accumulation of 10^6^ ions in the Orbitrap. The ten most intense peptide ions in each survey scan with an intensity above 2000 counts (to avoid triggering fragmentation too early during the peptide elution profile) and a charge state ≥ 2 were sequentially isolated at a target value of 10 000 and fragmented in the linear ion trap by collision-induced dissociation. Normalized collision energy was set to 35% with an activation time of 30 ms. Peaks selected for fragmentation were automatically put on a dynamic exclusion list for 30 s with a mass tolerance of ±10 ppm to avoid selecting the same ion for fragmentation more than once. The following parameters were used: the repeat count was set to 1, the exclusion list size limit was 500, singly charged precursors were rejected, and the maximum injection time was set at 500 and 300 ms for full MS and MS/MS scan events, respectively. For an optimal duty cycle, the fragment ion spectra were recorded in the LTQ mass spectrometer in parallel with the Orbitrap full scan detection.

For Orbitrap measurements, an external calibration was used before each injection series ensuring an overall error mass accuracy below 5 ppm for the detected peptides. MS data were saved in RAW file format (Thermo Fisher Scientific) using XCalibur 2.0.7 with tune 2.4. The data analysis was performed with Proline software 1.4 supported by Mascot Distiller and Mascot server (v2.5.1; http://www.matrixscience.com) database search engine for peptide and protein identification using its automatic decoy database search to calculate a false discovery rate (FDR) of 1% at the peptide level. MS/MS spectra were compared to the *Escherichia coli* Reference proteome set database containing the phage ribosomal proteins (UniProt release 2017_01, January 18 2017, 23022 sequences, 7070297 residues). Mass tolerance for MS and MS/MS was set at 10 ppm and 0.5 Da, respectively. The enzyme selectivity was set to full trypsin with one miscleavage allowed for samples S30AE and L7/L12 and the enzyme selectivity was set to full V8-DE with one miscleavage allowed for sample S21.

Protein modifications were fixed carbamidomethylation of cysteines, variable oxidation of methionine, variable acetylation of lysine, variable acetylation of N-terminal residues.

## ACKNOWLEDGEMENTS

This work was supported by grant ERC UE 340440 to PF and Agence Nationale pour la Recherche, Direction Générale de l’Armement (#ANR-14-ASTR-0001) to RG. CMM was supported by the European Molecular Biology Organization (ALTF 1562-2015) and Marie Curie Actions program from the European Commission (LTFCOFUND2013, GA-2013-609409); CG was supported by Direction Générale de l’Armement and Ministère de l’Enseignement supérieur et de la Recherche; SR and MBS were supported by Gordon and Betty Moore Foundation (#3305, 3790) and National Science Foundation (OCE#1536989) awards. The work conducted by the U.S. Department of Energy Joint Genome Institute, a DOE Office of Science User Facility, is supported under Contract No. DE-AC02-05CH11231. The Ohio Supercomputer is acknowledged for high-performance compute support. We thank Fanny Demay for the technical assistance and Sophie Chat for help with electron microscopy.

## Supporting information available

**Supplementary Table S1**: List and PFAM accessions of ribosomal protein domains searched for in viral proteomes.

**Supplementary Table S2**: Detection of ribosomal proteins in uncultivated viral genomes. Metagenomes were classified as “Environmental”, “Engineered”, or “Host-associated” according to the GOLD database (https://gold.jgi.doe.gov). Values of pN/pS were calculated for all ribosomal proteins in a contig covered > 10x and with at least 1 SNP detected.

**Supplementary Table S3**: List of 81 proteins identified in ribosomes purified from *E.coli* control cells, including detailed mass spectrometry information on peptide sequences.

**Supplementary Table S4**: List of 54 proteins identified in ribosomes purified from *E.coli* cells after expression of viral S21 protein, including detailed mass spectrometry information on peptide sequences. The table also includes extended information on phage protein identification in S21 ribosomal sample.

**Supplementary Table S5**: List of 80 proteins identified in ribosomes purified from *E.coli* cells after expression of viral L7/L12 protein, including detailed mass spectrometry information on peptide sequences. The table also includes extended information on phage protein identification in L7/L12 ribosomal sample.

**Supplementary Table S6**: List of 71 proteins identified in ribosomes purified from *E.coli* cells after expression of viral S30AE protein, including detailed mass spectrometry information on peptide sequenced. Beta-galactosidase was added as an internal control.

**Supplementary Table S7**: List of 279 proteins identified in crude *E.coli* cell extracts after expression of viral S30AE protein, including detailed mass spectrometry information on peptide sequenced.

